# A cross-species multimodal database of gene expression in cardiac hypertrophy

**DOI:** 10.1101/2025.01.27.635074

**Authors:** Christoph Sandmann, Jan D Lanzer, Frank Stein, Ellen Malovrh, Nicholas Rüdinger, Norbert Frey, Julio Saez-Rodriguez, Mirko Völkers

## Abstract

Changes in gene expression determine the pathophysiological response of the heart during increased workload and biomechanical stress.^1,2^ Understanding the complex gene expression regulation during physiological and pathological stress provides insight into mechanisms that mediate the transition from cardiac hypertrophy to heart failure. We curated and generated a multi-omics database of cardiac hypertrophy and heart failure and exemplified its usage by evaluating the regulation of UCK2, a gene that belongs to a conserved cardiomyocyte hypertrophy gene expression signature with unknown function in heart disease. To provide the community with simple access to the database, we developed CHEERIO (Cardiac HypErtrophy gEne expRessIOn Database), a free and easy-to-use interactive web application to explore rodent and human gene expression during early, late, physiological and pathological cardiac hypertrophy, as well as heart failure at transcriptome, translatome, proteome, bulk and single-cell resolution. CHEERIO is freely available at https://voelkerslab.shinyapps.io/CHEERIO/.

## Main

Controlled by a network of signaling cascades, the heart undergoes cellular remodeling in response to increased workload and biomechanical stress.^1–4^ While certain stimuli induce cardiac growth with preserved function, other forms of cardiac hypertrophy are associated with impaired function and cardiovascular events.^1,2,5,6^ Understanding the changes in cardiac gene expression and pathway activities is critical to identify molecular events that determine whether the heart will undergo physiological growth with preserved function or pathological growth with increased risk for cardiovascular mortality.

With the increasing complexity and volume of datasets, data-driven hypothesis generation is often challenged by the accessibility of research data.^7^ To support cardiovascular researchers in interrogating cardiac disease datasets, facilitating hypothesis exploration, and identifying novel disease signatures, robust data engineering efforts are essential.^7^ Currently, databases such as ReHeaT (Reference of the HEArt failure Transcriptome)^8,9^ provide comprehensive overview of gene expression patterns in heart failure, while e.g. TACOMA (Transverse Aortic COnstriction Multi-omics Analysis)^10^ enables gene- and transcript-level exploration of multi-omics data from murine transverse aortic constriction (TAC) models. However, these databases lack either multimodality, a primary focus on cardiac hypertrophy or human data while covering single cell/nucleus and bulk resolution. Additionally, they do not enable users to perform custom data comparisons or generate signatures tailored to specific research hypotheses.

We herein introduce the CHEERIO (Cardiac HypErtrophy gEne expRessIOn) database, which enables a comprehensive analysis of the gene expression response to physiological and pathological stress (Figure 1A). Covering bulk, single-cell RNA-sequencing (RNA-Seq), ribosome profiling (Ribo-Seq) as well as proteomic datasets of newly generated and publicly available cardiac hypertrophy data from rodents and humans, CHEERIO serves as a comprehensive platform to examine gene regulation during the cellular growth response of the heart. By combining the database with common gene expression signatures of heart failure and the fetal human heart, CHEERIO further reveals hypertrophy-associated gene regulation that may be caused primarily by cardiac failure rather than growth and identifies genes that belong to the fetal gene program. CHEERIO is available as a free and easy-to-use web application.

**Figure 1:**
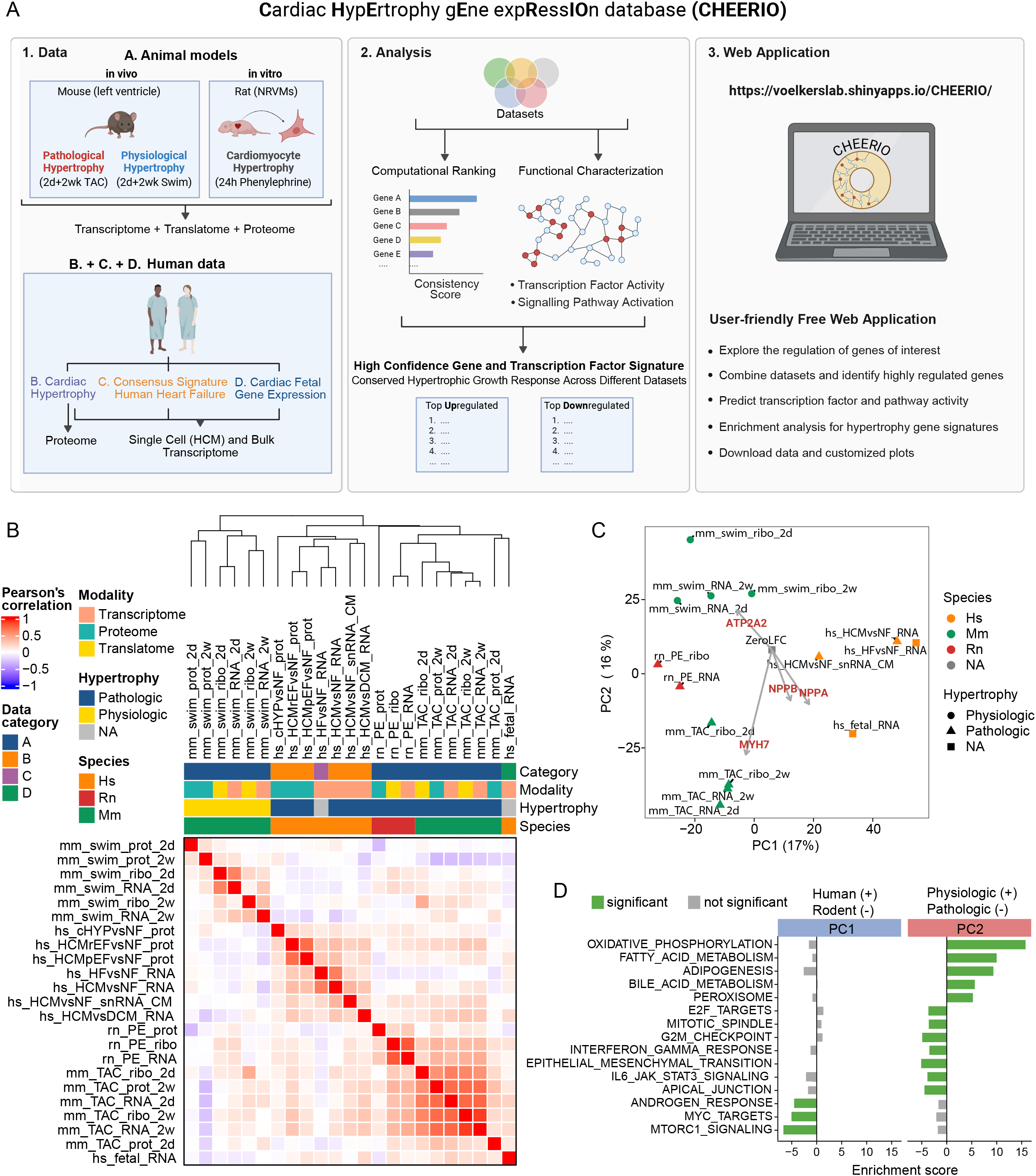
Integrated analysis of human and rodent cardiac hypertrophy gene expression datasets. **A**, Diagram displaying the experimental strategy used to derive the Cardiac Hypertrophy Gene Expression Database (CHEERIO). Transcriptome (bulk and single nuclear RNA-Seq), translatome (*in vitro*: bulk; and *in vivo*: cardiomyocyte-specific; Ribo-Seq), and proteome (bulk mass spectrometry) datasets of rodent and human cardiac hypertrophy were analyzed and integrated. A full list of all datasets included can be found in Table 1. Datasets are divided into the categories A. Animal models, B. Human Cardiac Hypertrophy (HCM), C. Human Heart Failure (DCM+ICM samples) and D. Human Fetal Gene Program. Animal models included *in vitro*: neonatal rat cardiomyocytes 24h phenylephrine treatment; and *in vivo*: 2d and 2wk transverse aortic constriction surgery for pathological hypertrophy or 2d and 2wk of swimming exercise for physiological hypertrophy. The human mass spectrometry data divides human samples into non failing hearts (cHyp) and failing hearts with reduced or preserved ejection fraction (HCM). All samples were compared to healthy controls. The fetal gene program compares healthy human fetal hearts to adult heart samples. Created in BioRender. **B**, Heatmap displaying Pearson’s correlaton of pariwise contrast comparisons. Categorical column annotations describe data category, modality, type of hypertrophy and species. **C**, Principal component analysis after exclusion of proteome data with first two components displayed. A log fold change vector of zero is added for interpretability with loadings of NPPA, NPPB, MYH7, ATP2A2 scaled by 10^3^ plotted starting from the zero log fold change vector. **D**, Gene set enrichment analysis on gene loadings for prinicpial components 1 (left) and 2 (right) with gene sets from MSIG DB. Green bars represent adjusted p-value < 0.05. hs: human; mm: mouse; rn: rat; RNA: RNA-seq (transcriptional regulation); snRNA: single nucleus RNA-Seq; ribo: Ribo-seq (translational regulation); prot: Mass spectrometry; 2d: two days; 2wk: two weeks; swim: swimming (physiologic hypertrophy); TAC: transverse-aortic-constriction (pathologic hypertrophy); PE: Phenylephrin; HCM: hypertrophic cardiomyopathy; HCMrEF: hypertrophic cardiomyopathy with reduced ejection fraction; HCMpEF: hypertrophic cardiomyopathy with preserved ejection fraction; cHYP: compensated cardiac hypertrophy (non-failing); DCM: dilated cardiomyopathy; NF: non-failing healthy heart.

We integrated 16 previously published gene expression datasets, which we supplemented with novel datasets of mouse proteomics after TAC surgery, as well as transcriptomics, translatomics and proteomics of mouse physiological hypertrophy after swimming exercise (Table 1). The datasets underwent an extensive quality control (Extended Data Fig. 1-7). Additionally, public datasets were curated from the literature for sufficient sample size and publicly available data at study initiation^8,11–16^. To avoid difficulties in comparing data from varying conditions, we limited the animal studies to a standardized set of data of identical timepoints and methods (24h neurohumoral stimulation with phenylephrine (PE) neonatal rat ventricular cardiomyocytes in vitro, 2d and 2wk TAC or Swim in 9-to-10-week-old C57BL/6N mice with RNA-Seq, Ribo-Seq and Mass spectrometry data). Methods are described in detail in the supplementary data. In total 1,439 samples were analyzed. Overall, we processed samples from rodent (n=74) and human (n=1,365) gene expression datasets of early, late, physiological, and pathological cardiac hypertrophy, as well as heart failure at transcriptome, translatome, proteome, bulk and single-cell resolution (Table 1). All samples were compared to respective healthy controls.

To facilitate cross-species and multi-modal analysis, we mapped genes and proteins to the human gene symbol taxonomy (HGNC) and retained only one-to-one mappings, which constituted 89% of mouse-to-human gene, 91% of mouse protein-to-gene, and 93% of rat-to-human gene conversions (Extended Data Fig. 6A). This resulted in 27,303 mapped protein-coding and noncoding genes across datasets (Extended Data Fig. 6B and C). After mapping, we compared datasets at the contrast level, leveraging case-control contrasts to minimize study-specific batch effects by avoiding direct gene expression comparisons across studies. For each group comparison, we independently calculated expression changes (log fold changes and p-values, see methods), which we will term contrasts. The number of differentially expressed genes varied between contrasts (range: 0–8,212; Q1: 5.75, median: 398, Q3: 2,056) (Extended Data Fig. 6D). Notably, proteomic analyses of mice after swim training did not identify differentially expressed proteins, suggesting that training effects at the selected timepoints are primarily restricted to RNA-level changes. The number of differentially expressed genes can be influenced by technical factors such as coverage and study design. To determine whether contrasts, despite variations in the number of differentially expressed genes, capture common trends of expression changes based on contrast characteristics such as data modality, species, and hypertrophy, we performed unsupervised analysis. Pairwise Pearson correlations were used to quantify the similarity between contrasts. Hierarchical clustering based on these correlation distances revealed distinct patterns of co-clustering, including a clear separation between pathologic and physiological hypertrophy datasets (Figure 1B), suggesting the capture of a fundamentally different gene expression program partially conserved across species.^1^ However, we observed species-specific clustering within the pathological hypertrophy group (Figure 1B), highlighting some limitations of rodent models for translational conclusions to humans, as previously noted by others.^17^ While this global comparison was limited to an intersection of 884 genes, we subsequently excluded proteomics datasets and compared RNA-Seq and Ribo-Seq contrasts across 4,493 genes. This analysis yielded a similar clustering pattern among the contrasts (Extended Data Fig. 7A), suggesting that this structure also generalizes to a higher number of genes. To ensure that this separation aligns with expected biological processes, we performed principal component analysis (PCA) after scaling logFC vectors to address their heteroscedastic distributions (Extended Data Fig. 7B and 7C). PCs 1 and 2 accounted for 33% of the total variance (Extended Data Fig. 7F), with PC1 separating species and PC2 differentiating physiological from pathological hypertrophy (Figure 1C). Gene set enrichment analysis of PC2 gene loadings confirmed known pathways that differentiate physiological and pathological hypertrophy, such as fatty acid oxidation and oxidative phosphorylation, inflammation and endothelial to mesenchymal transition/fibrosis (Figure 1D). In contrast, PC1 aligned to few signaling pathways, notably mTORC1, Myc and androgen signaling (Figure 1D), all of which are well established regulators of the cardiac growth response in mice. Their alignment suggests that these responses are more pronounced in mouse models compared to the human samples studied. In conclusion, contrasts derived from independent datasets reliably captured shared patterns of gene expression changes. These patterns align with known biological processes, reinforcing the distinction between physiological and pathological hypertrophy and highlighting the translational challenges of species-specific responses.

This compendium of gene expression changes can be leveraged to extract common gene signatures of contrasts of interest. To exemplify this, we selected contrasts to identify a conserved pathological cardiomyocyte hypertrophy disease signature that would be consistently regulated across modalities, models and species (Figure 2A-C). We combined transcriptomic and translatomic datasets (proteomic datasets were excluded due to limited gene coverage) of animal and human pathological cardiomyocyte hypertrophy and integrated p-values with a fisher combined test. From the top 500 significantly regulated genes (Supplementary Table 1) 272 agreed on directionality in at least 75% of datasets. Of those, 117 significantly regulated genes agreed on the direction of regulation across all datasets. These were selected for a conserved cardiomyocyte hypertrophy gene regulation signature, of which 67 are upregulated and 49 are downregulated during hypertrophic growth (Figure 2A-C, Supplementary Table 2). Among this conserved gene signature were known players regulated in heart disease, including NPPA, NPPB, ACTA1, XIRP2, MYH6 and ATP2A2 (Figure 2B).^1,2^ The conserved gene signature also included many genes recently identified in cardiomyocyte hypertrophy gene expression screens and functional analysis, such as PFKP^18^, RPL3^19^, or HSPB1^20^ among others. In addition, we identified numerous consistently regulated genes with unknown roles in cardiac hypertrophy, including RTN4, P3H2, RBP1, AGPAT4, EXT1, ECH1, HSDL2, RETSAT, LRP4 and UCK2, which may be of potential relevance for disease and therapy and are of high interest for future studies.

**Figure 2:**
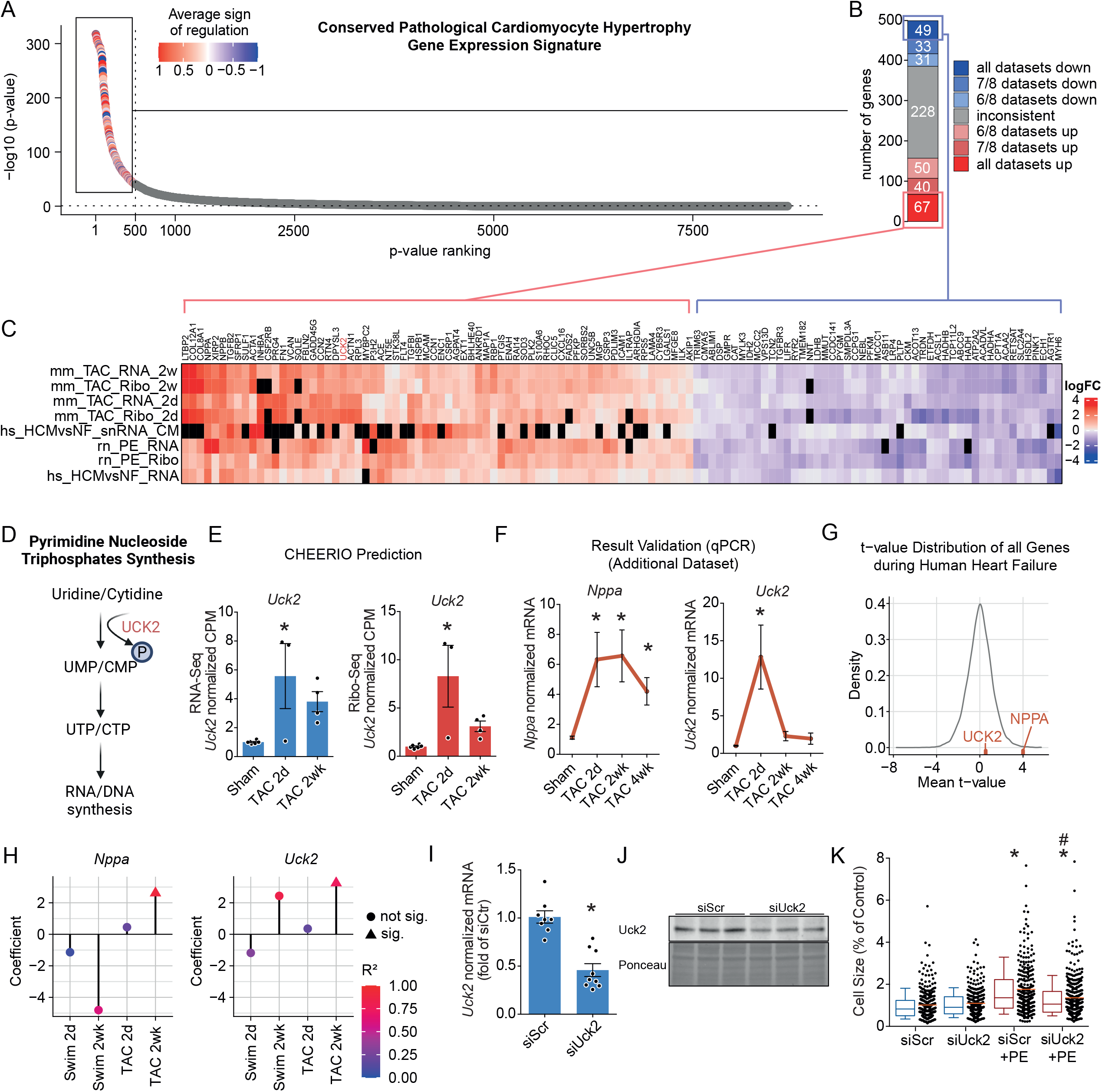
A consensus gene expression landscape of pathological hypertrophy identifies conserved gene regulation essential to cardiomyocyte growth. **A**, Gene ranking of conservation of gene regulation events in pathological cardiomyocyte hypertrophy across selected contrasts based on the adjusted p-value of a Fisher combined test. Selected contrasts included are rn_PE_RNA, rn_PE_Ribo, mm_TAC_RNA_2d, mm_TAC_RNA_2wk, mm_TAC_Ribo_2d, mm_TAC_Ribo_2wk, hs_HCMvsNF_RNA and hs_HCMvsNF_snRNA_CM. Proteomic datasets were excluded due to limited gene coverage. Vertical dotted line shows indicates top-500 genes which are colored by their mean sign regulation, and horizontal dotted line indicates p-value = 0.01. **B**, Fraction of genes among the top-500 regulated pathological cardiomyocyte hypertrophy genes with identical sign of regulation. Genes which were not regulated in the same direction in at least 6 of 8 contrasts were defined as inconsistent. At least 6 of 8 contrasts had to report a gene to be included in the donwstream analysis. **C**, Heatmap showing the log2 fold change regulation of the 67 top upregulated and 49 top downregulated pathological cardiomyocyte hypertrophy genes with consistent direction of regulation across all contrasts. Black squares indicate that the gene was not detected in that contrast. **D**, Diagram displaying the rate limiting enzymatic step of UCK2 in the synthesis of pyrimidine nucleoside triphosphates for RNA and DNA synthesis. Created in BioRender. **E**, Regulation of *UCK2* in mouse TAC RNA-Seq (blue) and Ribo-Seq (red) contrasts within CHEERIO. **F**, *Nppa* and *Uck2* mRNA levels in left ventricular lysates of an additional validation cohort of mice 2d, 2wk and 4wk after TAC surgery. Sham (merged 2d, 2wk and 4wk) n=18, TAC 2d n=3, TAC 2wk n=5, TAC 4wk n=7. **G**, Bulk trancriptomic t-value distribution of all genes during human heart failure from ReHeaT [cite] (DCM and ICM compared to healthy controls) with UCK2 and NPPA t-values labeled. **H**, Univariate linear models of the association between *Nppa* and *Uck2* gene expression and normalized mouse heart weight (heart weight/body weight). The coefficient of gene expression is displayed on the y-axis for different animal models (x-axis). Triangles indicate corrected p-value of the coefficient <0.05. The color indicates the R^2^ of the model. **I** and **J**, Normalized *Uck2* mRNA levels (I) and UCK2 protein levels (J) in neonatal rat ventricular cardiomyocytes after siRNA-mediated knock-down. **K**, Cross sectional surface area of neonatal rat ventricular cardiomyocytes after siRNA-mediated knock-down for 48h, followed by 50 μM PE treatment for 24h. Box plots show 10^th^ to 90^th^ percentile. Dot plots indicate individual cells. Approximately 300-400 cells were quantified per condition. * indicates p<0.05 from control, # indication p<0.05 from siScr+PE. hs: human; mm: mouse; rn: rat; RNA: RNA-seq (transcriptional regulation); snRNA: single nucleus RNA-Seq; ribo: Ribo-seq (translational regulation); 2d: two days; 2wk: two weeks; TAC: transverse-aortic-constriction (pathologic hypertrophy); PE: Phenylephrin; HCM: hypertrophic cardiomyopathy; NF: non-failing healthy heart.

To highlight the value of custom gene signatures generated by CHEERIO in identifying novel core genes of the cardiomyocyte growth response, we investigated UCK2, a consistently upregulated gene within the conserved hypertrophy signature whose role in cardiac hypertrophy remains unexplored. UCK2 encodes a pyrimidine ribonucleoside kinase which catalyzes the first step in the synthesis of pyrimidine nucleoside triphosphates required for RNA and DNA synthesis (Figure 2D). CHEERIO predicted upregulation of UCK2 at the transcriptome and translatome level (not detected in proteome datasets), which we confirmed in an independent cohort of mice after TAC surgery (Figure 2E-F). Contrary to many other genes of the conserved hypertrophy gene signature, such as NPPA, UCK2 is not regulated in heart failure as assessed in bulk transcriptomics (Figure 2G). UCK2 expression levels correlated with heart weight to body weight ratios in both physiological and pathological mouse cardiac hypertrophy (Fig. 2H), indicating that its upregulation is associated with heart growth in response to increased workload. In line with this was the observation that UCK2 also showed a trend for upregulation in all mice physiological hypertrophy datasets, which however only reached statistical significance within the 2wk Swim Ribo-Seq dataset. Importantly, UCK2 knockdown almost completely abolished hypertrophic growth of isolated cardiomyocytes in response to PE stimulation (Fig. 2I-K), highlighting that UCK2 is required for cardiomyocyte hypertrophy.

Next, we illustrated the use of CHEERIO to study the regulation of a particular gene of interest. We propose that consulting the expression profile of a gene across datasets in CHEERIO can help to refine a hypothesis and provide additional evidence to encourage an experimental investigation. We selected a novel Ensembl-annotated, conserved, and cardiac-enriched microprotein, SMIM4 (gene name recently renamed to UQCC5, previously also annotated as C3orf78) (Extended Data Fig. 8A-C), to ensure currently unknown regulation and involvement in cardiac pathophysiology. SMIM4 gene expression is moderately downregulated on the transcriptional, translational and protein level during cardiac hypertrophy, which we confirmed in an independent set of mouse hearts after TAC surgery in vivo (Extended Data Fig. 8D-E). Smim4 knockdown was not associated with altered cardiomyocyte hypertrophic growth in response to PE-stimulation, but significantly impaired cardiomyocyte metabolic activity as measured by an MTT assay, consistent with its recently described function as a respiratory chain assembly factor (Extended Data Fig. 8F-I). Thus, the regulation of SMIM4 may be involved in the metabolic switch that occurs during heart failure. Using CHEERIO, we were able to correctly depict the hitherto unknown regulation of a new microprotein encoding gene during cardiac hypertrophy and which showcases that CHEERIO may be used by other researchers to explore or confirm the regulation of genes of interest to save cost, time and the use of animals in their own studies.

To allow other researchers to easily access and make use of CHEERIO, we developed an interactive web application freely available at https://voelkerslab.shinyapps.io/CHEERIO/. The CHEERIO web application provides several key functions to evaluate gene expression and to estimate underlying transcription factor and signaling pathway activity, thus supporting researchers in elaborating, testing, and refining their hypothesis on the molecular nature of cardiac hypertrophy (Figure 6A). The CHEERIO web application is divided into several functional modules with simple user input and easy-to-understand visualizations and documentation.

First, users may identify how their genes of interest are regulated during cardiac hypertrophy and whether there exist differences between gene regulation at the transcriptional, translational or protein level, or between physiological and pathological hypertrophy (Figure 3A). Gene regulation of the genes of interest across the different datasets are displayed in the sections “A. Animal Models”, “B. Human Cardiac Hypertrophy”, “C. Human Heart Failure” and “D. Fetal Gene Program”. In each section, additional useful information is provided. For example, for the selected genes, known cardiovascular phenotypes of mouse models analyzed by the International Mouse Phenotyping Consortium are displayed, or the cell type-specific regulation in human hypertrophic cardiomyopathy is shown. As human cardiac hypertrophy samples are commonly derived from failing hearts of HCM patients, users are also informed whether their selected genes are similarly regulated in DCM and ICM heart failure patients as potential indication that gene regulation may not be specific to cardiac hypertrophy and may be influenced by the heart failure phenotype.

**Figure 3:**
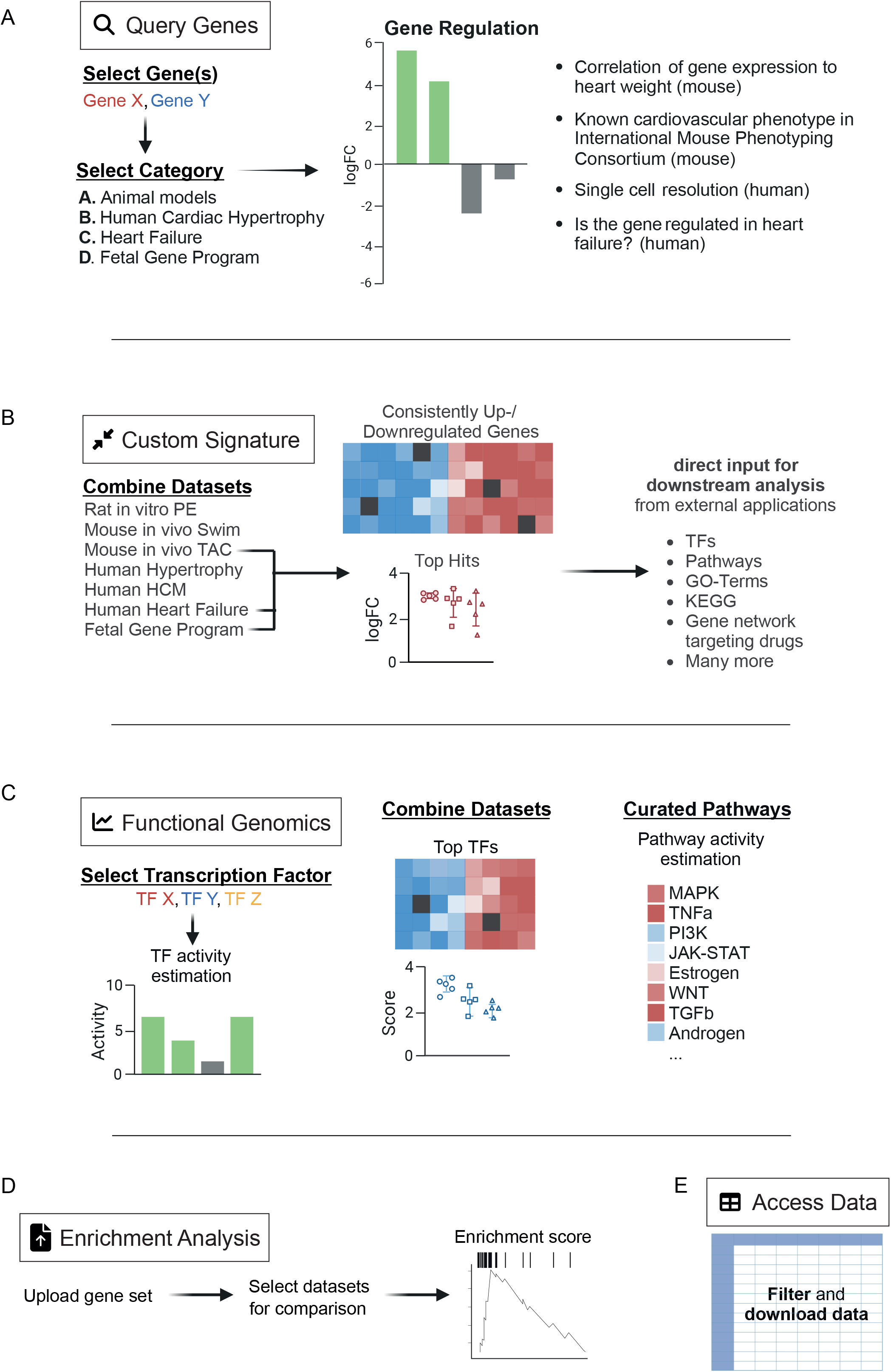
CHEERIO: a database and interactive web application for the analysis of gene expression in cardiac hypertrophy. CHEERIO is an easy-to-use web application for the exploration of gene regulation in cardiac hypertrophy available at https://voelkerslab.shinyapps.io/CHEERIO/. It consists of multiple functional modules: **A**, <u>Query Genes</u>: Users can search for genes of interest and explore their gene regulation in multiple contrast, supplemented with additional relevant information. **B**, <u>Custom Signature</u>: Users can combine datasets based on individual preferences and obtain top-regulated genes that can be used as a direct input to functional downstream analysis. **C**, <u>Functional Genomics</u>: Transcription factor and signaling pathway activity estimation can be derived for individual contrast. Users can either search for specific transcription factors or combine datasets to obtain top estimated TFs. **D**, <u>Enrichment Analysis</u>: Users can upload a gene set and test for its enrichment among cardiac hypertrophy datasets. **E**, <u>Access Data</u>: Data can be easily filtered and downloaded. Figure 3 created in BioRender.

Next, users may combine contrasts of their choice to identify genes highly regulated across the individually defined datasets to generate a custom disease signature. For this, users may select their desired datasets in the “Custom Signature” section, define an FDR cutoff, the numbers of genes to be plotted and the number of datasets in which a particular gene cannot be expressed before it is filtered out from the final list (Figure 3B). Several useful graphs are automatically plotted to give an overview of the selected analysis, such as the genes that show the strongest up- or downregulation across the selected datasets and their individual regulation in each condition. Gene lists of the top up- or downregulated genes with information on the regulation in each selected dataset can be downloaded. In addition, top signature genes can be copied to the clipboard and used as direct input to several useful applications that we cross-link within CHEERIO, allowing users to explore the output easily further for enrichment of signaling pathways, GO-terms, molecular signatures, KEGG-pathways, gene network-based drug interactions, tissue expression, related disorders, and others.

Next, using the section “Functional Genomics”, users may predict the activity of transcription factors of interest, potentially mediating transcriptional gene expression signatures observed across the various datasets (Figure 3C). For this, collecTRI^21^, a collection of gene regulatory networks, is used to predict transcription factor activities in each dataset. Similar to the “Gene Query” and “Custom Signature” section, users can either search for specific transcription factors they are interested in or combine datasets to identify transcription factors with shared activity patterns throughout the selected datasets. Further, PROGENy^22^, a resource that leverages a large compendium of publicly available signaling perturbation experiments to yield a common core of pathway responsive genes for human and mouse, is used to predict signaling pathway activities in each dataset.

n the “Enrichment Analysis” tab, users may upload a custom gene set to test for an enrichment in CHEERIO data sets (Fig. 3D). This enables users to corroborate their findings simultaneously in a number of related reference data sets and to explore the leading edge of the gene set consisting of genes driving a possible enrichment. We demonstrated this feature by enriching human plasma proteome signatures associated with increased or decreased risk of heart failure. We found that increased risk associated Plasama Proteins were enrichted in TAC signatures and identified IGFBP2, MFAP4, GDF15, FSTL1 as top marker driving this enrichment, suggesting that those markers might be, among others, of cardiac origin. These results can be reproduced by enabling the example data button selecting animal models for enrichment in the web app.

Finally, gene expression across every dataset of interest can be easily queried in the “Download Center”, to allow further in-depth analysis (Fig. 3E). Before downloading CSV or Excel lists of the data sets of interest, data can be further sorted or filtered if needed, while full data sets are available at zenodo https://doi.org/10.5281/zenodo.14509260.

The development of a phenotype is defined by cellular gene expression, controlled at various levels, including transcriptional, translational and post-translational regulation. Understanding how gene expression is regulated during cardiac disease is essential to identify underlying mechanisms that drive disease progression. To gain insights into the underlying gene expression signature of cardiac hypertrophy and its regulation, we analyzed various high-quality transcriptome, translatome and proteome datasets of rodent and human cardiac hypertrophy, including bulk and cell-type-specific data. Integrating those datasets, we identify a conserved pathological cardiomyocyte hypertrophy gene expression response that confirms known players in hypertrophic growth, but additionally uncovers many genes, currently uncharacterized in cardiac disease, that are highly consistently regulated in different rodent and human datasets and could serve as therapeutic targets of pathological cardiac hypertrophy.

In summary, here we present a large-scale integrated data analysis utilizing multiple highquality datasets on the regulation of gene expression at the transcriptional, translational, and proteomic levels in cardiac hypertrophy in rodents and humans, revealing underlying gene regulation of physiological and pathological hypertrophy. We extend the analysis with an easy-to-use interactive web application for data exploration, freely available at https://voelkerslab.shinyapps.io/CHEERIO/. We envision CHEERIO as a platform to inform users on gene expression regulation in cardiac hypertrophy and heart failure. Due to its modular design, CHEERIO can be easily extended. We plan to add more datasets in the future, including an increase in replicates, timepoints, disease entities, single cell level data and cardiac hypertrophy model systems.

## Acknowledgements

CS acknowledges the German Cardiac Society, the German Centre for Cardiovascular Research and the Dr. Rolf M. Schwiete Foundation. MV is supported by the DFG Heisenberg programme. NF, JSR and MV acknowledge the German Centre for Cardiovascular Research–Partner Site Heidelberg/Mannheim. NF, JSR, and MV were supported by Collaborative Research Center 1550 (CRC1550/SFB1550) “Molecular Circuits of Heart Disease.”

## Authors contributions

CS, JDL and MV conceived and designed the study. CS, EM and NR performed the experiments. CS, JDL and FS analyzed the data. JDL designed and implemented the web platform. CS, JSR and MV acquired funding. JSR and MV supervised the study. CH and JDL visualized the data; CH and JDL contributed to writing—original draft; CH, JDL, NF, JSR and MV contributed to writing—review and editing. All authors approved the final version of the article.

