## Supplementary material for "A cross-species multimodal database of gene expression in cardiac hypertrophy": Supplemantal data

### Methods

All animal experiments were approved by the institutional animal care and use committee of Heidelberg University. Data, analytical methods, and study materials are available from the corresponding author on reasonable request. Full datasets and code are available at zenodo <https://doi.org/10.5281/zenodo.14509260> and <https://github.com/saezlab/cheerio>.

**Preprocessing**

Datasets were individually processed, and differential gene expression analysis was performed with edgeR and limma R-packages. Processed heart failure studies and the fetal differential gene expression results were used from a published meta-analysis^1^. Processed single nuclear differential gene expression results were obtained from the original publication. Animal model and the Magnet data was reprocessed vie edgeR and limma. For gene conversions the biomaRt R-package^2^ was used to identify Ensembl gene ortholog annotations from mouse and rat to human gene symbols, one-to-one mappings were quantified and kept.

**Integrative Analysis**

Log2 fold-change (logFC) vectors were compared using Pearson’s correlation. The resulting correlation coefficients were transformed into a distance metric (1−coefficient)(1 - \text{coefficient})(1−coefficient) for hierarchical clustering using average linkage. Principal component analysis (PCA) was performed on contrast vectors excluding proteome datasets for increased coverage after scaling the logFC values. To enhance interpretability, a zero logFC vector was included in the PCA. In the PCA biplot, known hypertrophy markers were plotted, with their feature loadings scaled by a factor of 10³. PCA loadings were further analyzed for enrichment of hallmark gene sets from the MSigDB database^3^ using the run_ulm() function from the decoupleR R package^4^. A cross-species cardiac hypertrophy signature was generated by integrating logFC vectors from seven datasets. Fisher’s combined p-value was calculated for genes reported in at least 5 out of 7 contrasts, with p-values corrected using the Benjamini-Hochberg (BH) method. From this analysis, the top 500 genes were identified based on the elbow point of the -log10-transformed Fisher’s p-value ranking. To assess regulatory consistency, the number of contrasts reporting positive or negative logFC values was calculated. The analysis focused on 117 genes that were consistently regulated across contrasts and demonstrated the highest confidence.

**Web application**

We deployed an interactive database via the shiny R-package. The application includes the following analytical modules:

*Query Genes*

Users can filter selected genes across datasets to display their log-fold change, color-coded by a false discovery rate (FDR) threshold of <0.05. For animal models, univariate linear models were fitted with normalized heart weight as the dependent and gene expression as the independent variable. From these models, the gene expression coefficient, Benjamini-Hochberg (BH)-corrected p-value, and model R² were extracted and displayed. Additionally, phenotype associations for selected genes using SOLR query snytax from the International Mouse Phenotyping Consortium database.

*Custom Signature*

Selected contrasts are filtered by a user-defined FDR cutoff. Genes passing the filter in the specified number of contrasts selected by the user are retained and compared for fold-change consistency. Within each contrast, all genes were ranked based on their normalized signed, -log10-transformed p-value. This ranking is scaled between 0 (most upregulated) and 1 (most downregulated). For the custom signature a ranking is then derived by averaging these normalized ranks across contrasts. Top-ranked upregulated and downregulated genes are displayed for further analysis. These genes can be exported via a custom table, directly linked to *Drugstone*, or copied for enrichment analysis using *Enrichr* .

*Functional Genomics*

Transcription factor (TF) activities were inferred using the run_ulm() function from the *decoupleR* R package^4^, with gene regulatory networks provided by *collecTRI*^5^. A ranking statistic was computed by multiplying log-fold changes by their corresponding -log10-transformed p-values. This statistic was ranked and normalized relative to the gene vector length. Conserved TFs were ranked based on the mean activity values across contrasts. Pathway activities were computed using the top 500 pathway footprint genes from *PROGENy*^6^, analyzed with run_ulm() from the decoupleR R package^4^.

*Enrichment Analysis*

Users can perform enrichment analyses on uploaded gene sets. A gene statistic is calculated as the product of the -log10-transformed p-value and log-fold change. Gene set enrichment on this statistic is performed using the *fgsea* R package^7^.

**Laboratory animals**

All experiments were performed in 9- to 10-week-old male C57BL/6N mice. All animals were fed ad libitum and were housed at Heidelberg University in a temperature- and humidity-controlled facility with a 12-h light-dark cycle. Cardiomyocyte-specific Ribo-tag mice were previously described.^8^ Briefly, Ribo-tag mice (JAX ID 011029) were bred to αMHC-Cre mice to obtain Rpl22-HA-expressing homozygous mice in cardiomyocytes.

**Transverse aortic constriction surgery**

Animals were randomly assigned to the experimental groups. TAC was performed as previously described. Briefly, adult male mice were anesthetized using a 2% isofluorane/O2 mixture and intubated. Mice were treated with buprenorphine (0.1 mg/kg i.p.) and a partial trans-sternal thoracotomy performed using aseptic technique. An approximately 1.5 cm vertical left parasternal skin incision was made, underlying pectoralis muscle retracted, and the chest cavity entered through the fourth intercostal space. Using hooked retractors, adjacent ribs were retracted to expose the heart and aortic arch. The aorta was isolated from annexed tissue, and the artery partially ligated between the innominate and left common carotid arteries with 6-0 silk. The calibrated constriction of the aorta was performed by placing a dull 27- gauge needle to the side of the artery, the ligature tied firmly to both the needle and the artery, and, subsequently, the needle was removed leaving a calibrated stenosis of the aorta. Sham operated mice were exposed to the same procedure, except that the aorta was not constricted. The thoracic cavity was closed, and the animals were allowed to recover.

**Swimming exercise training**

Swimming exercise training was performed in a water tank for 90 minutes twice daily for 2 days or 2 weeks as previously described.^9^

**Preparation of tissue lysates**
Mice were sacrificed, and left ventricles were rapidly excised, washed in PBS and snap frozen in liquid nitrogen. Tissue used for Ribo-Seq and RNA-Seq analysis was washed in PBS containing 100 μg/ml cycloheximide. Left ventricles were homogenized using a tissue homogenizer in 5 volumes of ice-cold polysome buffer containing 20mM Tris pH 7.4, 10mM MgCl, 200mM KCl and 1% Trition X-100. Tissue used for Ribo-Seq and RNA-Seq analysis was homogenized in polysome buffer containing 20 mM Tris pH 7.4, 10 mM MgCl, 200 mM KCl, 2 mM DTT, 1% Triton X-100, 1U DNAse/μl and 100 μg/ml CHX. For library construction lysates were processed as previously described. For protein analysis via immunoblotting, initial lysates were further diluted with 9 volumes of RIPA buffer containing 20mM Tris-HCl (pH 7.4), 150mM NaCl, 1% Triton X-100, 0.1% SDS, 0,5% Sodium deoxycholate, protease inhibitor cOmplete ULTRA (cat# 05892791001, Roche) and phosphatase inhibitor PhosSTOP**‎** (cat# 04906837001, Sigma-Aldrich).

**Parallel generation of Ribo-seq and RNA-seq libraries**

Libraries were generated as previously described.^10^ For each animal, the heart was lysed in 700µl polysome buffer (20 mM Tris pH 7.4, 10 mM MgCl, 200 mM KCl, 2 mM DTT, 1% Triton X-100, 1U DNAse/μl and 100 μg/ml CHX) using a tissue homogenizer (Bullet Blender, NextAdvance). The tissue was homogenized further by passing the lysate through a 23-gauge syringe needle ten times. Homogenates were centrifuged at 4°C and 18,000xg for 10 min, and the supernatant was immediately used in the further steps. For complete lysis, the samples were kept on ice for 10 min and subsequently centrifuged at 20,000xg to precipitate cell debris. To accurately dissect translation and transcription, both Ribo-seq and RNA-seq libraries were prepared for each biological replicate from the identical lysate. Ribosome footprints were generated after immunoprecipitation of cardiomyocyte-specific polysomes with Anti-HA magnetic beads after treating the lysate with RNAse I (Ambion). Libraries were generated according to the mammalian Ribo-seq kit (Illumina). Barcodes were used to perform multiplex sequencing and create sequencing pools containing at least eight different samples and always an equal amount of both RNA and ribosome protected fragments (RPF) libraries. Sample pools were sequenced on the HiSeq 2000 platform using 50-bp sequencing chemistry.

**Sequencing data processing and quality control**

Sequencing data processing and quality control was performed as previously described.^2^ Adapters removal was done with Flexbar v3.0.3^6^ using standard filtering parameters (no prior trimming). Reads with more than 1 uncalled base were not included in the output: flexbar --may-uncalled 1 --pre-trim-left 0. Reads aligning to a custom bowtie2 v2.3.0^7^ index including mouse rRNA, mtRNA, tRNA and snRNA (Ensembl mus muculus release 97 – ncrna) were discarded. Remaining reads were then aligned in genomic coordinates to the mouse genome (GRCm38.p6) with STAR, v.2.5.3a^8^, inserting annotations on the fly: STAR --quantMode TranscriptomeSAM --alignIntronMin 20 --alignIntronMax 100000 --outFilterMismatchNmax 1 --outFilterIntronMotifs RemoveNoncanonicalUnannotated --outFilterMismatchNoverLmax 0.04 --sjdbOverhang 50. Only uniquely mapping reads were kept for analysis. For Ribo-seq data, only periodic fragment lengths were kept that showed a distinctive triplet periodicity. We used the automatic Bayesian selection of read lengths and ribosome P-site offsets (BPPS) method^9^ to select and shift aligned reads to properly account for P-site of the ribosome. For the RNA-seq data, reads were trimmed from the 3' end after adapter removal, such that the read length before alignment did match the maximum periodic fragment length of the corresponding Ribo-seq sample, as determined with the BPPS method. Finally, abundance estimates and read count to coding sequences were obtained using HTSeq-count^11^, taking into account the strand-specific protocols. We used an FDR<0.01 as a cutoff.

**Gene Ontology analysis**

For GO term analysis, genes with FDR <0.05 were considered for further analysis. GOTERM_BP_DIRECT in DAVID 2021^12^ was used with the subset of expressed protein-coding genes as background set. Only enriched GO terms with at least five significantly changed genes were kept for further analysis and ranked by p-value. Top enriched terms were retained and visualized with a custom plotting routine showing p-value of enrichment.

**Mass spectrometry sample preparation SP3 and TMT labeling, OASIS**

Reduction of disulfide bridges in cysteine containing proteins was performed with dithiothreitol (56°C, 30 min, 10 mM in 50 mM HEPES, pH 8.5). Reduced cysteines were alkylated with 2-chloroacetamide (room temperature, in the dark, 30 min, 20 mM in 50 mM HEPES, pH 8.5). Samples were prepared using the SP3 protocol^13^ and trypsin (sequencing grade, Promega) was added in an enzyme to protein ratio 1:50 for overnight digestion at 37°C. Next day, peptide recovery in HEPES buffer by collecting supernatant on magnet and combining with second elution wash of beads with HEPES buffer. Peptides were labelled with TMT10plex^14^ Isobaric Label Reagent (ThermoFisher) according to the manufacturer’s instructions. Samples were combined for the TMT10plex and for further sample clean up an OASIS® HLB µElution Plate (Waters) was used. Offline high pH reverse phase fractionation was carried out on an Agilent 1200 Infinity high-performance liquid chromatography system, equipped with a Gemini C18 column (3 μm, 110 Å, 100 x 1.0 mm, Phenomenex).

**LC-MS/MS**

An UltiMate 3000 RSLC nano LC system (Dionex) fitted with a trapping cartridge (µ-Precolumn C18 PepMap 100, 5µm, 300 µm i.d. x 5 mm, 100 Å) and an analytical column (nanoEase™ M/Z HSS T3 column 75 µm x 250 mm C18, 1.8 µm, 100 Å, Waters). Trapping was carried out with a constant flow of 0.05% trifluoroacetic acid at 30 µL/min onto the trapping column for 6 minutes. Subsequently, peptides were eluted via the analytical column with a constant flow of 0.3 µL/min with increasing percentage of solvent B (0.1% formic acid in acetonitrile). The outlet of the analytical column was coupled directly to a QExactive plus (Thermo) mass spectrometer using the Nanospray Flex™ ion source in positive ion mode. The peptides were introduced into the QExactive plus via a Pico-Tip Emitter 360 µm OD x 20 µm ID; 10 µm tip (New Objective) and an applied spray voltage of 2.3 kV. The capillary temperature was set at 320°C. Full mass scan was acquired with mass range 375-1200 m/z in profile mode with resolution of 70000. The filling time was set at maximum of 10 ms with a limitation of 3x10^6^ ions. Data dependent acquisition (DDA) was performed with the resolution of the Orbitrap set to 35000, with a fill time of 120 ms and a limitation of 2x10^5^ ions. A normalized collision energy of 32 was applied. Dynamic exclusion time of 30 s was used. The peptide match algorithm was set to ‘preferred’ and charge exclusion ‘unassigned’, charge states 1, 5 - 8 were excluded. MS data was acquired in profile mode.

**Mass spectrometry data analysis**

IsobarQuant^15^ and Mascot (v2.2.07) were used to process the acquired data, which was searched against a Mus musculus GRCm38 pep database containing common contaminants and reversed sequences. The following modifications were included into the search parameters: Carbamidomethyl (C) and TMT10 (K) (fixed modification), Acetyl (Protein N-term), Oxidation (M) and TMT10 (N-term) (variable modifications). For the full scan (MS1) a mass error tolerance of 10 ppm and for MS/MS (MS2) spectra of 0.02 Da was set. Further parameters were set: Trypsin as protease with an allowance of maximum two missed cleavages: a minimum peptide length of seven amino acids; at least two unique peptides were required for a protein identification. The false discovery rate on peptide and protein level was set to 0.01. The raw output files of IsobarQuant (protein.txt – files) were processed using the R programming language (ISBN 3-900051-07-0). Only proteins that were quantified with at least two unique peptides were considered for the analysis. Raw TMT intensities (signal_sum columns) were first cleaned for batch effects using limma^16^ and further normalized using vsn (variance stabilization normalization)^17^. Proteins were tested for differential expression using the limma package. The replicate information was added as a factor in the design matrix given as an argument to the ‘lmFit’ function of limma. A protein was annotated as a hit with a false discovery rate (fdr) smaller 5 % and a fold-change of at least 50 % and as a candidate with a fdr below 25 % and a fold-change of at least 50 %.

**Cultured Cardiomyocytes**

Neonatal rat ventricular cardiomyocytes were isolated from one to two-day old animals via enzymatic digestion and purified by Percoll density gradient centrifugation as previously described.^10^ Briefly, cardiomyocytes were plated at a density of 0.4 - 0.5 X 10^6^ cells per well on 22.6mm or 34.8 mm well plastic plates that had been pre-treated with 0.1% gelatine for 1 h at 37 °C (cat# F1141, Sigma-Aldrich) and then cultured in DMEM/F12 1:1 (cat# 11330032, Thermo Fisher Scientific), containing 10% fetal bovine serum, 100 units/mL of penicillin, 100µg/mL streptomycin and 292 μg/ml glutamine (cat# 10378016, Thermo Fisher Scientific). After 24h media was changed to DMEM/F12 1:1 supplemented with 0.5% fetal bovine serum, 100 units/mL of penicillin, 100µg/mL streptomycin and 292 μg/ml glutamine. After an additional 24 hours, cells were treated with Dulbecco´s phosphate buffered saline (PBS) (cat# 14190-144, Thermo Fisher Scientific) as a control or (R)-(-)-phenylephrine hydrochloride (50µM) (cat# P6126, Sigma-Aldrich) diluted in PBS, for 24h

**RNA interference**

Pre‐designed synthetic anti‐rat Uck2 small interfering RNA (s156426), anti-rat Smim4 siRNA (s166843), and scrambled siRNA (Silencer™ Select Negative Control No. 1 siRNA) as negative control were purchased from Thermo Fisher Scientific. Isolated neonatal cardiomyocytes were transfected with 25 nM final concentration of siRNAs using the HiPerfect transfection reagent as previously described (cat# 301704, QIAGEN).^18^

**Cultured cell area**

Phase-contrast microscopy pictures were taken from five different fields per culture (n=3 individual cell cultures for each treatment). Cell area was determined using FIJI. The cell area of at least 300 cells was measured for each condition.

**MTT assay**

7000 cells per well were plated in 96-well plates that had been pre-treated with 0.1% gelatine for 1 h at 37 °C (cat# F1141, Sigma-Aldrich) and then cultured as described above. After knockdown and PE treatment, MTT (Vybrant® MTT Cell Proliferation Assay Kit (V13154), Thermo Fisher Scientific) to the cells according to manufacturer instructions. After 4 hours 10% SDS with 0.01M HCl was added to the cells for solubilization. After overnight incubation, absorbance was measured at 570nm using an EnSpire® MultimodePlate Reader (PerkinElmer).

**Immunoblotting**

Cultured cells were lysed in RIPA Buffer consisting of 50mM Tris pH 7.5, 150mM NaCl, 1% Triton X-100 and 1% SDS, which was supplemented with protease inhibitor cOmplete ULTRA (cat# 05892791001, Roche) phosphatase inhibitor PhosSTOP**‎** (cat# 04906837001, Roche). Tissue lysates were prepared as described above. Lysates were cleared by centrifugation at 4 °C for 10 minutes at 20.000 rcf. Lysate protein concentration was determined using the DC Protein Assay Kit II (cat# 5000112, Bio-Rad) according to the manufacturer's instructions. Equivalent amounts of protein, usually 20-30 µg, were brought up to similar volume, mixed with Laemmli Sample Buffer (Bio-Rad; 161-0747) and 2-Mercapoethanol (cat# M6250, Sigma-Aldrich) and boiled at 95°C for 5 minutes. Samples were separated on SDS-PAGE gels and transferred to Immobilon-P transfer membranes (cat# IPVH00010, Merck Millipore). The following antibodies were used to probe the membranes: UCK2 (1:1000, cat# 10511-1-AP, Proteintech), SMIM4 (1:500, cat# BSS-BS-15183R, Biozol). Quantifications of all immunoblots were normalized to a loading control (Ponceau).

**Quantitative Real Time PCR** 
Total RNA was isolated from cultured cardiomyocytes using the Quick-RNA MiniPrep Kit (cat# R1055, Zymo Research) and from tissue using TRIzol (cat# 15596026, Invitrogen). cDNA was generated by reverse transcription using the High-Capacity cDNA Reverse Transcription Kit (cat# 4368814, Thermo Fisher). Quantitative Real Time PCR was performed with qPCR SYBR® Green Master Mix (cat# SL-9903R). The following primers were used:

Mouse-Uck2-Fw: AGGAGTTCTGCTTGCCAACAA

Mouse-Uck2-Rev: CCGTTGAGATAGCCGTTCGT

Rat-Uck2-Fw: TCGTCTCCCACTCACGGAAA

Rat-Uck2-Rev: CAGACGGGTATCTGCATCGG

Mouse-Smim4-Fw: AAGCAGCGATTCGGCATCTA

Mouse-Smim4-Rev: TCCAGCCTTCTCTGATACTGTCT

Rat-Smim4-Fw: TTTGTCCTGGGAGGAACGATG

Rat-Smim4-Rev: ATCTTCCAGCCTTCTCTGATACTG

Mouse-and-Rat-18S-Fw: CGAGCCGCCTGGATACC

Mouse-and-Rat-18S-Rev: CATGGCCTCAGTTCCGAAAA

**Statistical Analysis**

Statistical analysis for bar graphs or dot plots was performed using GraphPad Prism 7.0 (Graphpad Software Inc; www.graphpad.com) or R (R Foundation; https://www.r-project.org). Data values are mean ± standard error of the mean. For statistical analysis one-way ANOVA with Tukey post-hoc analysis was used. When only two conditions were compared, unpaired two tailed t-test was used. p < 0.05 was defined as a significant difference.

**Extended Data Figure Legends**

**Extended Data Figure 1: Quality control of Ribo-Seq data of mouse sham/TAC and sedentary/swim libraries.**

**A** and **B**, Ribo-seq read counts of all libraries showing periodic (usable), non-periodic, multi-mapped and non-aligning reads, when mapped to the mouse transcriptome for TAC (A) and swim (B) samples. **C** and **D**, Beeswarm plots visualizing the sequenced ribosome footprint lengths across all samples from TAC (C) and swim (D) datasets. **E** and **F**, Bar plot showing the percentage of reads mapping to the coding sequence (CDS) and 5′ and 3′ untranslated regions (UTR) of annotated protein-coding genes for TAC (E) and swim (F) samples. Each line represents a separate sample. **G** and **H**, Bar plot summarizing the ribosome protected footprint periodicity for all samples as the percentage of footprints that match the three reading frames of the annotated coding sequence genome wide for TAC (G) and swim samples (H). **I**, Graphical representation of the periodic profile Bayesian model selection showing results for different read lengths and p-site offsets for one typical library.

**Extended Data Figure 2: Gene expression quality control of Ribo-Seq data of mouse sham/TAC and sedentary/swim datasets.**

**A**, Heart weight to body weight ratios of mouse hearts after sham surgery and 2d or 2wk after TAC surgery used to generate the RNA-Seq and Ribo-Seq libraries of this study. **B** and **C**, Principal component analysis of RNA-seq (B) and Ribo-seq (C) libraries after sham or TAC surgery. **D**, Heart weight to body weight ratios of mouse hearts after sedentary or swimming exercise for 2d or 2wk used to generate the RNA-Seq and Ribo-Seq libraries of this study. **E** and **F**, Principal component analysis of RNA-seq (E) and Ribo-seq (F) libraries after sedentary or swimming exercise. **G**, Venn diagram of gene products detected in the transcriptome, translatome and proteome datasets of left ventricular lysates 2 days after TAC surgery used in this study. **H** and **I**, Scatterplots of Ribo-Seq vs RNA-seq in sham- and TAC-operated mice 2d (H) and 2wk (I) after surgery. Transcripts were considered significant when false discovery rate <0.01. Gray dots indicate no significant change. Significant change at transcriptional level is shown in blue, at translational level in red, and regulation at both translational and transcriptional levels in green. **J**, Venn diagram of gene products detected in the transcriptome, translatome and proteome datasets of left ventricular lysates 2 days after swimming exercise used in this study. **K** and **L**, Scatterplots of Ribo-Seq vs RNA-seq of mice after sedentary or swimming exercise for 2d (K) or 2wk (L). Transcripts were considered significant when false discovery rate <0.01. Gray dots indicate no significant change. Significant change at transcriptional level is shown in blue, at translational level in red, and regulation at both translational and transcriptional levels in green.

**Extended Data Figure 3: Pathological and physiological cardiac hypertrophy marker gene expression in mouse hearts after TAC surgery.**

Gene expression regulation in RNA-seq, Ribo-seq and mass spectrometry dataset of mouse left ventricular lysates after TAC surgery for the pathological cardiac hypertrophy marker genes (**A**) Nppa, (**B**) Nppb, (**C**) Myh6, (**D**) Myh7, (**E**) Col1a1, (**F**) Fn1, and the physiological cardiac hypertrophy marker genes (**G**) Pdk4 and (**H**) Ppargc1a.

**Extended Data Figure 4: Pathological and physiological cardiac hypertrophy marker gene expression in mouse hearts after swimming exercise.**

Gene expression regulation in RNA-seq, Ribo-seq and mass spectrometry dataset of mouse left ventricular lysates after swimming exercise for the pathological cardiac hypertrophy marker genes (**A**) Nppa, (**B**) Nppb, (**C**) Myh6, (**D**) Myh7, (**E**) Col1a1, (**F**) Fn1, and the physiological cardiac hypertrophy marker genes (**G**) Pdk4 and (**H**) Ppargc1a.

**Extended Data Figure 5: Enrichment of biological process in pathological and physiological mouse gene expression datasets.**

Enrichment of GO terms (biological process) of significantly regulated genes/transcripts/proteins in RNA-Seq (**A** and **B**), Ribo-Seq (**C** and **D**) and mass spectrometry (**E** and **F**) data of left ventricular lysates 2d or 2wk after TAC surgery and of RNA-Seq (**G** and **H**), Ribo-Seq (**I** and **J**) and mass spectrometry (**K** and **L**) data of left ventricular lysates 2d or 2wk after swimming exercise. Detected genes were used as the respective gene expression background of each dataset. The 10 most significant GO terms by p-value per group are displayed. No GO terms were significantly enriched in the conditions of G, I, K and L.

**Extended Data Figure 6:** **Overview of gene mapping and coverage.**

**A**, Gene orthology mapping between protein and gene names as well as between species. Only one-to-one mappings were retained for downstream analysis. **B**, Histogram displaying the number of genes (y-axis) that are captured by different number of contrasts (x-axis). **C**, Gene coverage after qualtiy control and filtering for all contrasts in CHEERIO. **D**, Overview of differentially expressed genes (DEGs) and their logFC reported by selected contrasts. Top panel displays number of DEGs at corrected p-value <0.05 reported by each contrast; Bottom panel displays log fold change distribution with points colored by significance (upregulated in red, downregulated in blue, nonsignificant in grey) and with contrast labes (x-axis) colored by data category.

**Extended Data Figure 7: Additional contrast comparisons for a subset excluding proteome data.**

**A**, Heatmap displaying Pearson’s correlation between selected contrasts excluding proteomic data sets (same contrasts selected as for the PCA in figure 2C). **B** and **C**,  Distribution of log fold change values for contrasts for PCA analysis. Heteroscedasticity was accounted for by scaling (C). **D**, Screeplot displaying explained variance (%) for principial components.

**Extended Figure 8: Gene expression regulation and functional evaluation of the microprotein Smim4 in cardiac hypertrophy.**

**A**, Diagram showing the strategy to identify heart-enriched microproteins with evidence of translation in humans, used to identify an example gene with unknown regulation in cardiac hypertrophy. 798 protein coding transcripts were heart-enriched in the evo-devo database of mammalian organs from Cardosa-Moreira et al. Of those, 260 were microproteins with Ensembl annotation, thus potentially included in CHEERIO. 253 of 260 Ensembl annotated microprotein transcripts showed evidence of translation in human hearts (van Heesch et al.). Among those we chose SMIM4 for further evaluation within CHEERIO. **B**, *SMIM4* transcript levels throughout prenatal and postnatal development in human, macaque, mouse and rabbit brain, heart, kidney, liver and testis. Dotted lines indicate birth. **C**, Coding sequence conservation of Smim4 in humans, mice and rats. **D**, Regulation of *Smim4* in mouse TAC RNA-Seq (blue), Ribo-Seq (red) and mass spectrometry (yellow) contrasts within CHEERIO. **E**, *Smim4* mRNA levels in left ventricular lysates of an additional validation cohort of mice 2d, 2wk and 4wk after TAC surgery. Sham (merged 2d, 2wk and 4wk) n=18, TAC 2d n=3, TAC 2wk n=5, TAC 4wk n=7. **F** and **G**, Normalized *Smim4* mRNA levels (F) and SMIM4 protein levels (G) in neonatal rat ventricular cardiomyocytes 48h after siRNA-mediated knock-down. **H**, Cross sectional surface area of neonatal rat ventricular cardiomyocytes after siRNA-mediated knock-down for 48h, followed by 50 μM PE treatment for 24h. Box plots show 10^th^ to 90^th^ percentile. Dot plots indicate individual cells. Approximately 300-400 cells were quantified per condition. **I**, Metabolic activity of neonatal rat ventricular cardiomyocytes quantified by MTT assay after siRNA-mediated knock-down and PE treatment. * indicates p<0.05 from control, # indication p<0.05 from siScr+PE.
